# Evolutionary trade-offs between growth and reproduction under obesogenic conditions: sex-biased skeletal and gonadal maturation in mice

**DOI:** 10.64898/2026.07.16.738527

**Authors:** Clarissa R. Coveney, Amar Sarkar, Neil G.O. Ibata, David E. Maridas, Devandar Kaushik, Alanah McKeever, Jenna Shoman, Seiji Aoki, Talia Kahan, Cary Allen-Blevins, Danalaxshmi S. Ganapathee, Alexander S. Okamoto, Gayani Senevirathne, Emily M. Venable, Grace Rubin, Laura D. Schell, Victoria A. Tobolsky, Daniel E. Lieberman, Rachel N. Carmody, Terence D. Capellini

## Abstract

Adolescence is a brief period during which finite energy resources are reallocated from linear growth toward reproductive maturation. Rising childhood obesity and earlier onset of puberty suggest that modern, energy rich diets may distort these evolved energy allocation rules, but causal mechanisms remain unclear. Here, we expose male and female wild-type and leptin receptor deficient (*Db/Db*) to either high fat or normal chow diets. We longitudinally analyze metabolic, skeletal, gonadal, endocrine and insulin-receptor phenotypes from 4-10 weeks of age (sexual maturation window in mice). In WT males, HFDs increased adiposity and impaired glucose tolerance, and selectively remodeled joint morphology, advanced gonadal maturation, and shifted insulin receptor expression from growth plates to testes. Together, these changes indicate a rebalancing of energy use toward reproduction at the expense of skeletal and metabolic health. WT females showed subtler systemic metabolic disruption but clear diet-responsive changes in growth plate and ovarian maturation. Our findings indicate that energy-rich diets during adolescence shift evolved allocation rules to prioritize reproductive readiness over skeletal robustness in a sex-dependent manner, with potential consequences for precocious puberty and bone health in humans.

## Introduction

Obesity rates amongst children and adolescents have been rising across the globe for the past four decades, especially in developed countries^1^. Though obesity exerts numerous adverse effects on the health^2^ and social lives of individuals^3^, it may also play a significant role in altering growth and developmental trajectories^4,5^. This concern is apparent during adolescence; the brief, energetically expensive window during which mammals achieve sexual maturity and terminal body size. For humans in Western countries, the average age of onset for puberty has fallen by 3 years over the past century^6^, and globally by 3 months per decade from 1977-2013^7^, a decline that is strongly correlated with increased prevalence of childhood obesity^7,8^. Similarly, there is also a significant positive correlation between body mass index (BMI) and pubertal onset^9^. These kinds of patterns suggest that body size and the timing of sexual maturation may be connected to one another. However, we possess limited knowledge of how altered energy supplies (e.g., under high fat or high sugar Western diets) impact cellular and tissue function across growth-related phenotypes (e.g., the skeleton) and reproduction- related phenotypes (e.g., the gonads) during adolescence.

With respect to the relationship between body size and sexual maturity, both epidemiological and *in vivo* mouse studies suggest that obesity does not simply add mass during adolescence. Rather, obesity can reshape the trajectory and quality of other organ systems, including skeletal maturation in ways that are sexually dimorphic and mechanistically linked to energy signaling. The transition to reproductive investment during adolescence is in large part mediated by endocrine signals reflecting nutritional and energy status. During human adolescence, individuals will build and accrue between a third to half of their skeleton that they will maintain for life, reaching peak bone mass at age 25^10^. Obesity has been shown to increase bone mass and accelerate growth with diminishing returns on bone health long term^11–13^. For example, overweight and obese children exhibit increased rates of long-bone fractures relative to their normal-weight peers, suggesting a mismatch between rapid gains in body size and the capacity of bone and joint tissues to maintain appropriate strength during this period^14^. In parallel, longitudinal cohorts indicate that higher childhood BMI is associated with greater height, which is consistent with the idea that energy-rich environments can accelerate both linear growth and sexual maturation^15^. Together, these findings raise concerns that obesity-driven acceleration of growth and sexual development may come at the cost of compromised skeletal growth, perturbed joint morphology^16^, and reduced skeletal robustness^12^ during a window when these tissues are normally being fine-tuned for lifelong use.

Of course, these correlational patterns preclude inferences of causality. Thus, it is important to investigate the associations between obesity, growth, and sexual development using experimental methods in the context of a theoretical framework that explicitly relates physical growth to sexual maturity. In the present work, we combine an experimental murine model with life history theory to integrate and investigate the relationships between obesity, growth, and reproduction. Life history theory is particularly useful because it frames adolescence as an evolutionary strategy that redirects limited energy supplies from growth toward reproduction^17,18^. Under this framework, attaining sexual maturity is a dynamic (rather than static) phenomenon that is dependent on energy availability and ecological risk^17,19^. Such research will help in understanding how the environment can affect individual development under changing environments such as the energy rich human nutritional landscape.

Here, using variations in diet and mouse genotype, we aim to elucidate with greater precision the mechanisms of global body energy allocation^17,19,20^. We hypothesized that an energetic surplus (high fat diet) during adolescence would redirect energy allocation away from skeletal growth and toward reproductive maturation earlier in sexual maturation. We anticipated a HFD to advance gonadal development potentially through altered insulin receptor upregulation in skeletal growth plates and gonads, with potential sex-specific patterns reflecting different evolutionary constraints on male and female life histories. By comparing WT and leptin receptor–deficient (Db/Db) mice, we further predicted that intact leptin signalling would be required to translate obesogenic diets into coordinated shifts across metabolic, skeletal and reproductive systems.

We investigate how the energetic surplus characteristic of modern diets interacts with this critical developmental window to reshape patterns of growth and maturation, and how these effects differ between males and females. We focus on wild-type (WT) and leptin receptor–deficient (*Db*/*Db*) mice exposed to either normal chow (NC) or a high-fat diet (HFD). We examine mice from 4 to 10 weeks of age, a period that encompasses rapid skeletal growth and reproductive maturity. Across this window, we combine longitudinal metabolic phenotyping with morphometric analysis of the hindlimb using high- resolution MicroCT analysis, histological assessment of growth plate and articular cartilage, quantitative analyses of gonadal histology, endocrine profiling, and tissue-level measures of insulin receptor expression in bone and gonadal tissues, in regulating metabolic and endocrine function. We test the hypothesis that obesogenic diets during the sexual maturation window selectively accelerate gonadal maturation and skeletal growth, enabling examinations of whether rapid gains in size are prioritized over reproductive readiness or vice versa. This integrative approach allows us to examine how energy-rich environments perturb evolved allocation dynamics that normally coordinate growth and reproduction, and to identify sex-specific pathways through which adolescent obesity may generate lasting risks for skeletal and reproductive function.

## Methods

### Ethics

All experiments performed on mice, including euthanasia, were approved by Harvard University’s Institutional Animal Care and Use Committees with protocol 13-04- 161-2. No human participants were used.

### Animals

Wild-type (WT, C57BL/6J, #000664, Jackson Labs) and leptin receptor deficient (B6.BKS(D)-*Lepr^db^*/J (*Db/Db*), #000697, Jackson Labs) mice were placed on either a high-fat diet or normal chow beginning at four weeks of age and aged to 10 weeks (minimum 7–8 mice per sex, diet, and genotype group; total n = 63). A further group of WT male and female mice were placed on a NC or HFD for 2 or 4 weeks (n =6 per group) to assess longitudinal changes in sexual maturation window.

### Diets

High fat diet (HFD) (Envigo TD.06414): 60.3% calories from fat, 21.4% calories from carbohydrate, 18.3% calories from protein. Normal chow (NC) diet (5P75 Prolab® IsoPro® RMH 3000): 14.3% calories from fat, 59.5% calories from carbohydrate, 26.1% calories from protein.

### Glucose tolerance tests and insulin sensitivity tests

Glucose tolerance tests and insulin sensitivity tests were conducted biweekly in parallel with body composition assessment by EchoMRI (assessed body fat%, lean mass% and water). For both glucose tolerance tests and insulin sensitivity tests, mice were fasted for approximately 4-6h and subsequently administered 1.5 mg glucose/g body mass or insulin-sensitivity tests, 1 U/kg body mass of insulin (Humulin R) by intraperitoneal injection. Blood glucose levels were measured from tail blood at 0, 15, 30, 60, 90, and 120 min after glucose administration using a Clarity BG1000 blood glucometer (Clarity Diagnostics) and test strips. All measurements were performed in duplicate, with additional replicates obtained when the difference between initial readings was ≥20 mg/dL.

### Plasma collection and plasma profiling

For end point plasma of 4- and 10-week-old mice, blood was collected from mice via cardiac puncture, yielding 0.5–0.75 mL of blood. Mice were euthanized, confirmed by cervical dislocation, followed by the opening of the thoracic cavity, exposing the heart. A 23-gauge needle attached to a 1 mL syringe was used to puncture the right ventricle and withdraw blood. Blood was transferred into potassium EDTA containing tubes, treated with DPPIV (Sigma, DPP4-M) and Pefabloc (Roche, AEBSF), centrifuged (2,500-3,000 × g for 10 min to obtain plasma). Longitudinal tail blood samples were collected at 4, 6 and 8 weeks of age for the longitudinal study.

All longitudinal plasma samples were stored at −80 °C until analysis by Luminex multiplex Steroid-Thyroid Hormone 6-Plex Discovery Assay (Eve Technologies). The 6- plex assay included Cortisol, Estradiol, Progesterone, T3, T4, and Testosterone. Plasma was assessed at a dilution of 1:2, in duplicate. All terminal plasma samples were stored at −80 °C until analysis Mouse Bone Metabolism 13-Plex Panel (Creative Proteomics). The 13-plex assay included ACTH, DKK-1, FGF-23, IL-1β, IL-6, Insulin, Leptin, PTH, OC, OPG, OPN, SOST, TNFα. All analytes were measured in duplicate.

### MicroCT

Post euthanasia, skin and excess muscle were removed from each hind limb before fixation in 10% neutral buffered formalin for 24 hours at room temperature. Left hind limbs were collected, formalin-fixed, and prepared for high-resolution MicroCT imaging in 70% ethanol. The acetabulum, femur and tibia from right hindlimbs were imaged by high-resolution micro–computed tomography (μCT40, SCANCO Medical AG). Scans were acquired at 12 μm isotropic voxel size, 70 kVp peak tube voltage, 114 μA tube current and 200 ms integration time. Digital Imaging and Communication in Medicine (DICOM) files were exported and analyzed in OsiriX MD v7.5 (Pixmeo SARL), and measurements were obtained following established anatomical protocols ^21,22^. Briefly, we quantified: acetabular depth, diameter and inclination; proximal femoral geometry (valgus cut angle, neck–shaft angle, neck length, neck diameter, head offset, head diameter); distal femoral morphology (bicondylar width, notch width, medial and lateral condylar widths and curvatures, medial/central/lateral trochlear widths, trochlear groove depth, trochlear angle); and proximal tibial morphology (plateau width, medial and lateral posterior tibial slope, and medial and lateral tibial spine height). Quantification of bone density metrics for the trabecular and cortical bone were made using SCANCO Medical AG software following published guidelines^23^.

### Histology

Following MicroCT, limbs were then decalcified in 14% EDTA at a pH 7.5 for eight days. Formalin- fixed tissues were sent to the Massachusetts General Hospital Center for Skeletal Research Histology and Histomorphometry Services core facility and processed for sectioning and histologic staining using fast green and Safranin O staining. Embedding, sectioning, and staining were performed without knowledge of genotype. A minimum of 6 coronal sections were taken throughout the middle of the joint (60–80-μm levels).

### Cartilage Thickness Measurements

Cartilage thickness, including of the non-calcified and calcified regions, was measured as previously described^24^, using ImageJ. Briefly, the mean of three measurements from the medial and lateral tibial plateaus using Safranin O and Fast Green stained coronal knee sections. Measurements were only taken from 3 consecutive sections in the middle of the joint for a total of 18 measurements per section. Similarly for growth plate length, measurements were taken (as previously described^25^) from the top of the tibial growth plate to the trabecular bone. Across the width of the tibial growth plate, 9 measurements were taken per histological section. Each joint had 5 or 6 histological sections, therefore each joint has 45 or 54 total growth plate measurements that are averaged to obtain the total measurement of the tibial growth plate for the joint.

### Cartilage integrity (OARSI) scoring

Safranin O stained coronal knee joint sections were assessed using the summed OARSI score as previously described^24^ in order to measure the cartilage health within the joint. The scale of OARSI scoring ranges from 0- 6, with 0 indicating healthy cartilage and 6 indicating degraded cartilage down to the subchondral bone. Both the lateral and medial tibial plateaus and the lateral and medial femoral condyles are assessed. Sections were blind scored by two readers (DK and SA) using the summed Osteoarthritis Research Society International (OARSI) scoring method of OA^26^.

### Immunohistochemistry insulin receptor

To stain for insulin receptor presence, unstained, fixed coronal knee joint sections were deparaffinized, rehydrated, then quenched in 0.3 M glycine for 40 minutes and treated with proteinase K for 30 minutes in a humidified chamber at 37 ℃. Sections were then treated with 0.1 unit chondroitinase for 30 minutes at 37 ℃. Sections are permeabilized in Triton X-100 for 15 minutes before they are blocked with 10% goat serum and 10% bovine serum albumin in phosphate-buffered saline. The cartilage sections are incubated overnight at 4℃ in either insulin receptor beta antibody (E9L5V, Cell Signaling) at 1:50, IgG control at a 1:50, or no primary antibody. After incubation, sections were washed and incubated with Alexa Fluor 555 secondary antibody at 1:500 for 1 hour. Sections were incubated in DAPI nuclear stain at 1:5000 before being mounted with ProLong Gold (P3693, Invitrogen) before being imaged using Zeiss Axioscan 7 and analyzed with Zeiss Zen Microscopy Software. For gonadal tissues, the protocol was the same, excluding the chondroitinase treatment.

### Histological analysis of gonadal tissues

Gonadal tissues were collected from 3, 6, and 8 week old WT male and female mice, fixed in 4% PFA before being embedded in paraffin and sectioned and stained with Hematoxylin and Eosin. Quantification of ovarian follicles and seminiferous tubules was performed using ImageJ software^27^. Briefly, for ovarian samples, follicular size was measured using cross-sectional area and diameter from histological sections, with diameter measured across the widest point of the follicle within a given section, with a minimum of 5 sections throughout the ovary used per animal. Seminiferous tubule diameter was measured from round or near-round transverse sections to avoid oblique distortion, as previously described^28^. A minimum of three measurements was taken per sample and averaged for analysis.

### Statistics

All analyses were conducted in R. Unless otherwise stated, statistical significance was set at p = 0.05, and p-values from multiple pairwise comparisons were adjusted using the Benjamini-Hochberg procedure to control the false discovery rate. For most endpoint comparisons between diet, sex and genotype, we used non-parametric Wilcoxon rank sum (Mann Whitney U) tests because group sizes were modest and several variables deviated from normality. Where longitudinal data were available (e.g. body mass, GTT/IST AUC, repeated hormone measures), we fitted linear or generalized linear mixed-effects models with mouse identity as a random intercept and fixed effects of sex, genotype, diet, age and their interactions where appropriate. For the MicroCT dataset, principal component analysis (PCA) was performed on morphometric variables to summarize multivariate variation in joint shape and bone geometry; PCs were interpreted based on trait loadings and compared across groups using non-parametric tests. Outliers were identified by visual inspection (boxplots, residual plots) and retained; missing values were left as missing and not imputed.

## Results

Wild-type (WT) and leptin receptor deficient (*Db/Db*) mice were placed on either a high-fat diet or normal chow beginning at four weeks of age and aged to 10 weeks. At baseline and every 2 weeks thereafter, we performed glucose tolerance tests (GTT), insulin sensitivity tests (IST), EchoMRI body composition measurements, and both longitudinal and endpoint sacrifice plasma collection. At sacrifice, gonadal and subcutaneous fat depots were also isolated.

Over the 6-week period, both sexes displayed increases in body mass. However, by 10 weeks, only the WT males in the HFD condition showed a significant increase in body weight relative to males in the NC condition (Fig. 1a). This effect was driven primarily by an increase in body-fat percentage and fat mass (Fig. 1b, and raw weights Supplementary Fig. 1a), without detectable reductions in lean-mass percentage or lean mass (Supplementary Fig. 1a). In *Db*/*Db* mice, both males and females gained substantial weight with age in both the NC and HFD conditions (Fig. 1a), accompanied by increases in both fat and lean mass (Fig. 1b and Supplementary Fig. 1a). WT males in the HFD condition showed significantly greater gonadal fat mass than WT males in the NC condition, whereas gonadal fat did not increase in WT females in the HFD condition (Fig. 1c and Supplementary Fig. 1b). In contrast, both male and female WT mice in the HFD condition showed subcutaneous fat increased significantly with HFD, with larger absolute gains in males (Fig. 1c and Supplementary Fig. 1b). Overall, *Db*/*Db* mice displayed markedly higher masses of both gonadal and subcutaneous fat than WT animals (Fig. 1c and Supplementary Fig. 1b). In *Db*/*Db* mice, HFD further increased both fat depots in females but exerted little additional effects in males, the opposite pattern to WT. By 10 weeks of age, WT males in the HFD condition exhibited impaired glucose tolerance without a detectable change in insulin sensitivity, whereas WT females showed no significant diet effect on either GTT or IST (Fig. 1d). In both male and female *Db*/*Db* mice, glucose tolerance and insulin sensitivity were already severely impaired and were exacerbated in the HFD condition, frequently exceeding the upper range of the glucometer (Fig. 1d). Differences in circulating insulin concentrations by diet, sex, or genotype did not reach statistical significance, but tended to be descriptively higher in WT mice of both sexes in the HFD condition (Fig. 1e). In contrast, leptin concentrations were significantly elevated only in WT males in the HFD condition and were substantially higher in *Db*/*Db* mice than in WT mice across diets and sexes (Supplementary Fig. 1c).

**Fig 1.**
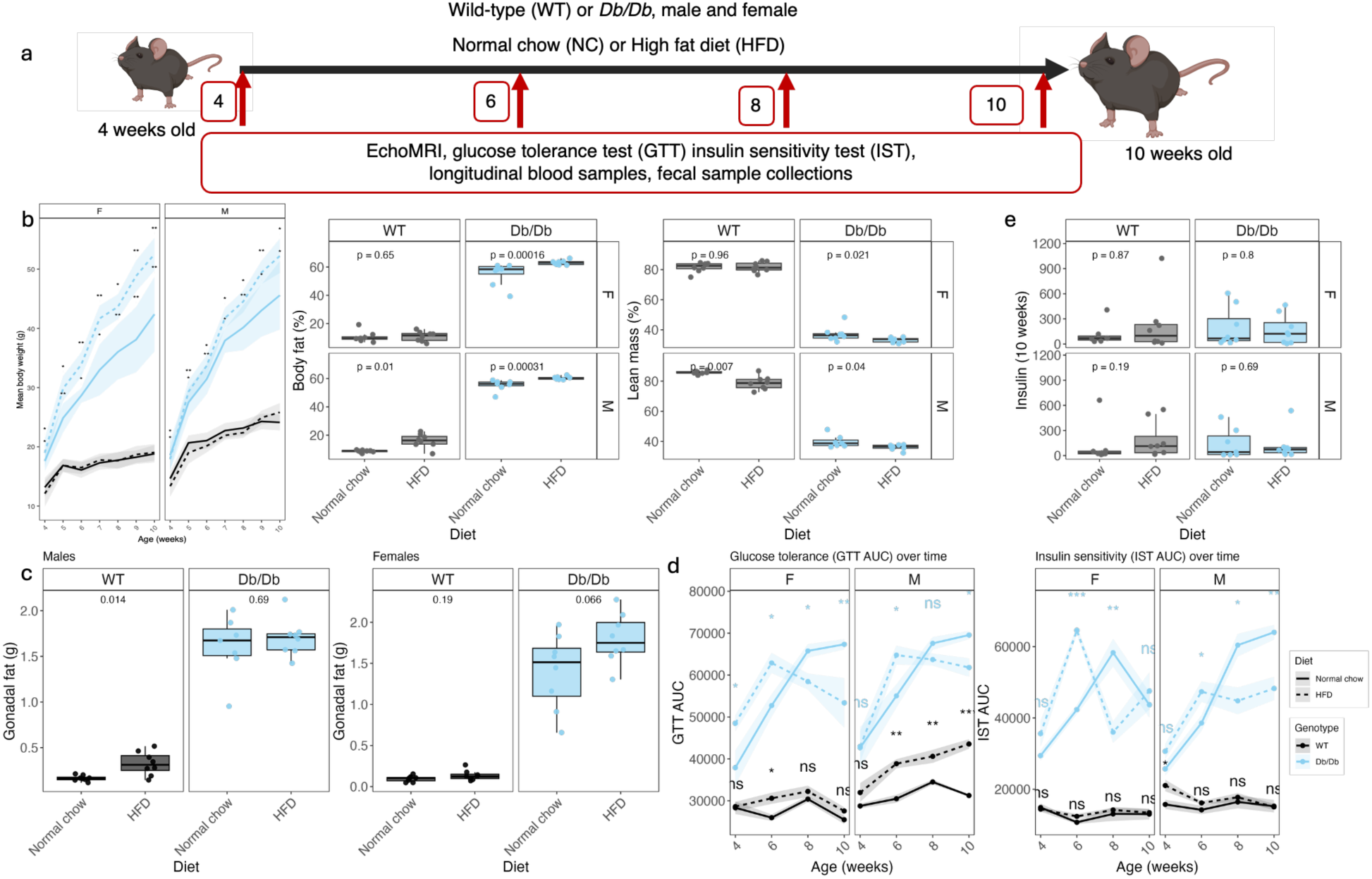
High fat diet and obesogenic models drive metabolic changes during sexual maturation in male, but not female, mice. **a)** Schematic overview of the experiment. **b)** Body fat and lean mass percentages calculated from EchoMRI measurements. **c)** Raw gonadal and subcut fat weights collected at sacrifice. **d)** Area under the curve (AUC) of glucose tolerance tests (GTT) and insulin sensitivity tests (IST) measured at 4, 6, 8 and 10 weeks in WT and *Db/Db* mice in the NC and HFD conditions. Lines and ribbons represent group mean ± SEM. **e)** Plasma concentration of insulin (pg/mL) at 10 weeks of age. Wilcoxon rank–sum test (Mann–Whitney U), (ns *p* ≥ 0.05, \**p* < 0.05, \*\**p* < 0.01, \*\*\**p* < 0.001). For each sex WT NC *n* = 8, WT HFD *n* = 8, *Db*/*Db* NC *n* = 7-8, *Db*/*Db* HFD *n* = 8.

We first used microCT of the right hindlimb to quantify 29 morphometric traits spanning distal femoral joint shape and diaphyseal length (Fig. 2a). Principal component analysis revealed marked separation of WT and *Db*/*Db* mice in both sexes (35.5% variance explained by PC1, 9.0% variance explained by PC2; Fig. 2b), with additional within-genotype shifts by diet (Supplementary Fig. 2a). This suggests that HFDs reshaped joint morphology and cortical bone density without altering linear growth: tibia length as well as overall body length (nose to base of tail), were comparable between NC and HFD conditions within each genotype and sex (Fig. 2c and Supplementary Fig. 2b). In contrast, specific joint surfaces were selectively remodeled: WT females in the HFD condition showed a reduction in medial trochlear width and an increase in medial tibial spine height relative to mice in the NC condition, whereas WT males displayed a trend toward narrower tibial width (Fig. 2c, Supplementary Fig. 2b, **Supplementary Table 1**).

**Figure 2.**
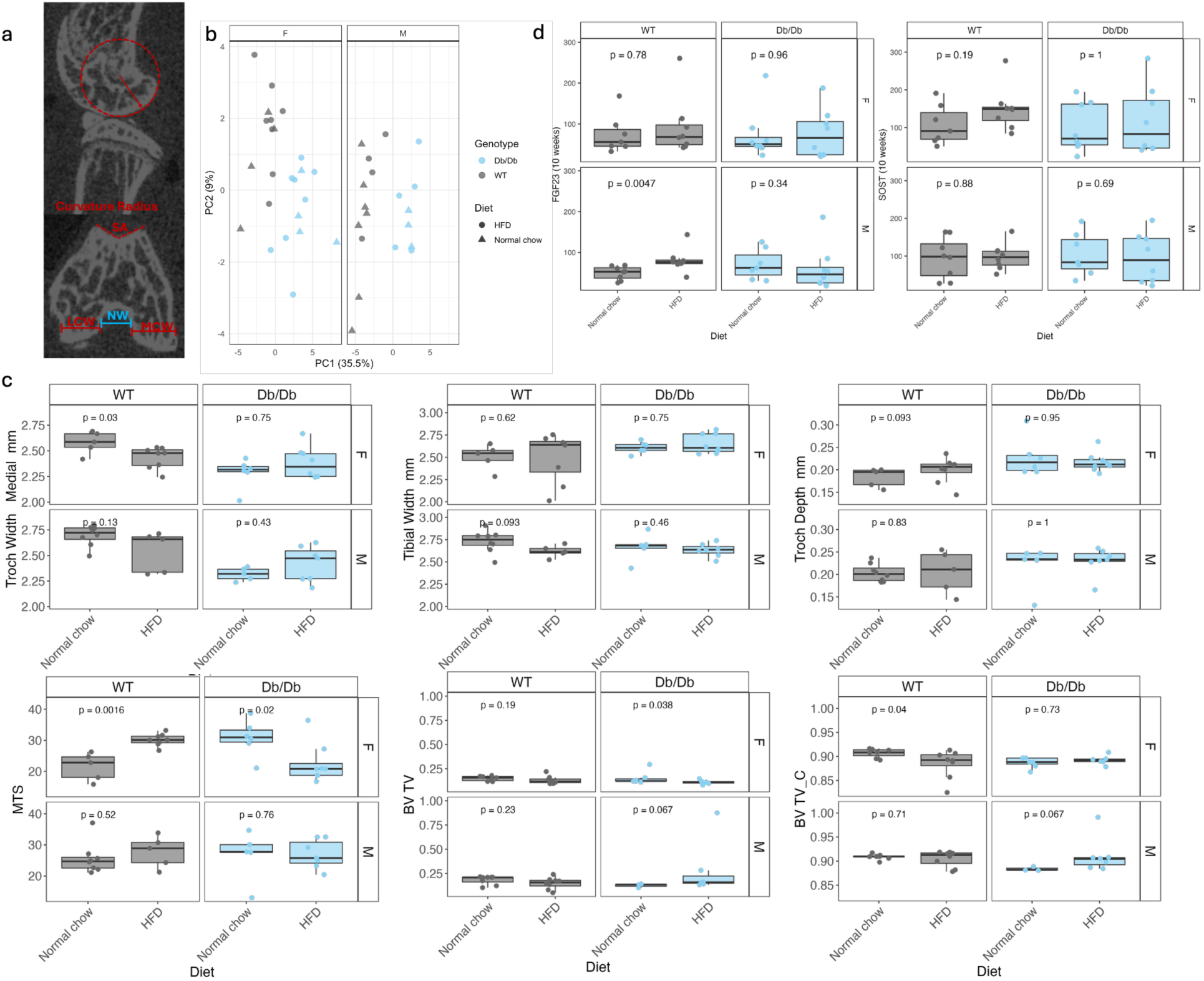
High fat diet and obesogenic models result in altered joint morphology and bone density but have little impact on bone turnover markers. **a)** Representative MicroCT images of measurements. **b)** PCA plots of all morphological measurements. **c)** Plots of measurements that showed statistically significant differences between diets for each sex and genotype including BV/TV for cortical bone. **d)** Bone turnover markers FGF23 and SOST. Symbols represent individual mice. Wilcoxon rank–sum test (Mann– Whitney U)). For each sex and genotype *n*= 5-7.

These effects of the diet were blunted or absent in *Db*/*Db* mice. Across genotypes, cortical bone density (BV/TV_C) was lower in HFD-fed mice, reaching statistical significance in females but not males (Fig. 2c). At the endocrine level, plasma FGF23 concentrations were higher in males than females at 10 weeks, whereas SOST levels did not differ significantly by genotype or diet (Fig. 2d). In general, HFDs did not produce differences either by sex or genotype in plasma concentrations of other bone turnover markers including, DKK1, osteocalcin (OC), osteopontin (OPN), parathyroid hormone (PTH), and osteoprotegerin (OPG) (however, male *Db*/*Db* mice displayed elevated levels of OPG). There were also no observed changes in ACTH or cortisol concentrations. Together, these data show that a high fat diet during sexual maturation and growth decouples overall size from cortical bone accrual and selectively remodels joint morphology in a sex dependent manner.

To determine whether HFDs alter articular cartilage or growth plate maturation, we performed histology on paraffin-embedded knee joints from the left hindlimb. In mice, articular cartilage progressively calcifies between 3 and 10 weeks of age as the tidemark advances^24^ (Fig. 3a, Supplementary Fig. 3a). To investigate whether HFDs impact skeletal cartilage maturation, we quantified the thickness of uncalcified cartilage (superficial to the tidemark; green line, Fig. 3a) and calcified cartilage (deep zone to the tidemark; blue line, Fig. 3a). Because previous work has not detected sex differences in cartilage thickness, these measurements were not separated by sex^24^. HFDs measurably affected neither the total articular cartilage thickness nor the relative thickness of uncalcified versus calcified zones, in either genotype or sex (Fig. 3b). Consistent with this result, quantitative assessment of cartilage integrity using the gold standard OARSI scoring method^29^ revealed no evidence of diet- or genotype-dependent degeneration at 10 weeks in WT mice, but showed elevated OARSI disease scores in Db/Db mice in the HFD condition (Supplementary Fig. 3b).

**Figure 3.**
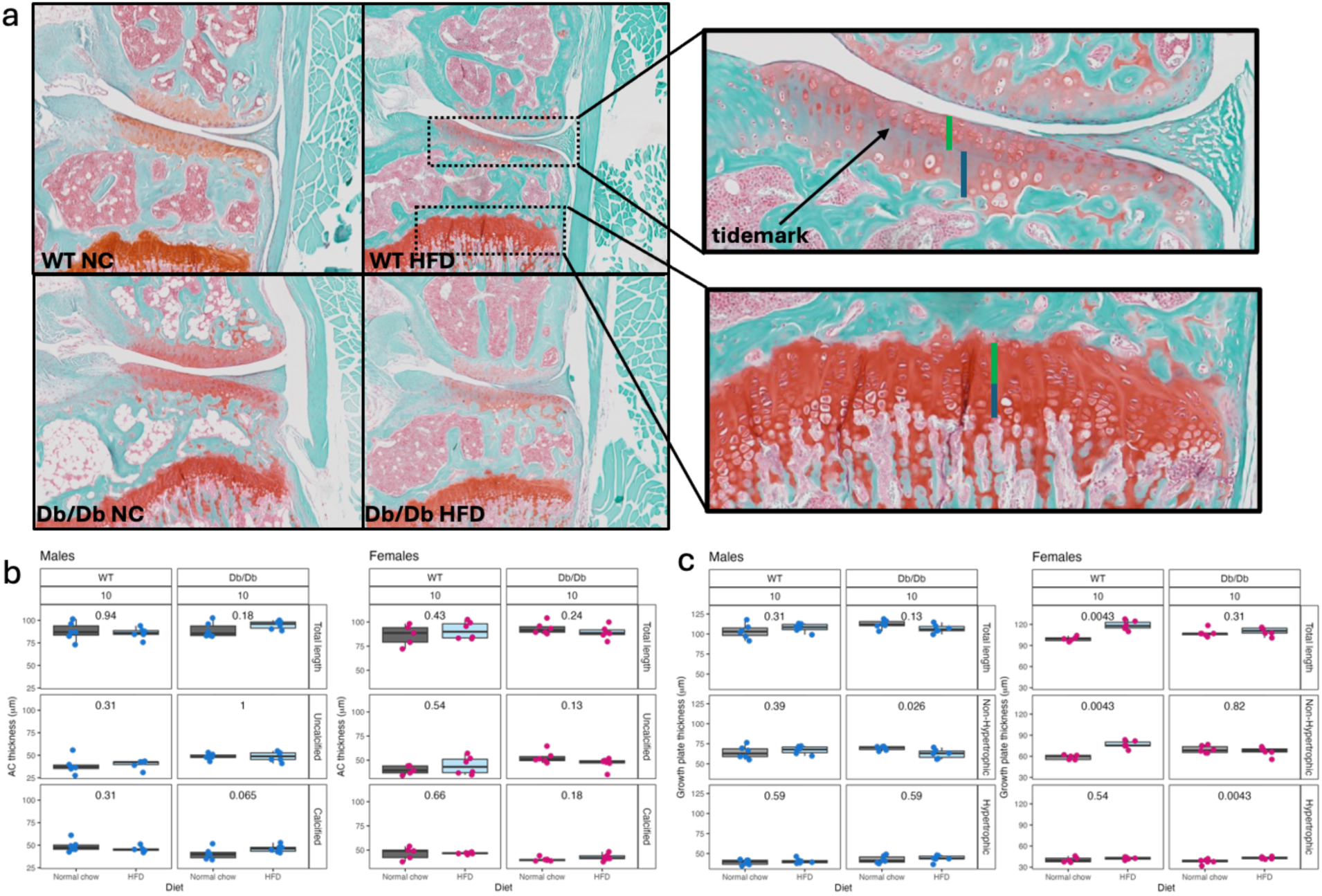
High fat diet alters growth plate chondrocyte populations in female mice but does not impact on articular cartilage maturation. **a)** Coronal histological sections of 10 week-old WT and *Db*/*Db* male and female mice were stained with Safranin O for histologic analysis. Medial compartment shown depicting uncalcified (green line) and calcified (blue line) articular and growth plate non-hypertrophic (green line) and hypertophic (blue line) cartilage. **b)** At 10 weeks of age, thickness of the articular cartilage as well as the uncalcified and calcified regions of both medial and lateral tibial plateaus were measured and averaged. **c)** At 10 weeks of age, total length of the growth plate cartilage as well as the non-hypertrophic and hypertrophic regions across the entire width of the tibia were taken and averaged. Symbols represent individual mice. Wilcoxon rank- sum test (Mann–Whitney *U*) For each sex and genotype, *n*= 5-8 in any group.

We next examined the growth plate cartilage, where chondrocytes rapidly divide in the proliferative zones, move into the resting zone, and then differentiate into hypertrophic chondrocytes before replacement by bone. We delineated the non-hypertrophic region (resting/proliferative zone; green line, Fig. 3a) and hypertrophic region (blue line, Fig. 3a) and measured their lengths as markers of growth plate elongation and maturation. In WT females, total growth plate length was significantly greater in the HFD condition animals than in the NC condition, driven entirely by an expansion of the non-hypertrophic region (Fig. 3c). WT males showed no detectable dietary effect on growth plate chondrocyte proportions (Fig. 3c). In *Db*/*Db* females, HFDs increased the length of the hypertrophic zone, without a corresponding change in the non-hypertrophic region or total length (Fig. 3c), whilst *Db*/*Db* males experienced no changes to overall length but had shorter hypertrophic regions.

We next examined how HFDs during this period affect maturation of gonadal tissues and plasma hormones. We performed histology on the ovaries and testes (Fig. 4a and Supplementary Fig. 4a) from 3-, 6-, and, 8-week old WT mice in the NC or HFD conditions, spanning the interval when mice reach sexual maturity (females can reproduce from ∼8 weeks of age). In the ovaries, we quantified follicle length as a read-out of follicular growth and maturation. During the developmental process, mouse ovaries are initially highly active, rapidly developing follicles for later reproduction (Supplementary Figure 4a). We averaged follicle length (Fig. 4b and Supplementary Fig. 4b) per animal and compared 6 and 8 weeks of age measurements (6-week NC, 6-week HFD, 8-week NC, 8-week HFD). In 6-week old mice in the NC condition, the ovaries had significantly larger follicles than ovaries from 6-week old mice in the HFD condition, 8-week old mice in the NC condition, and 8-week old mice in the HFD condition (adjusted *p ≤* 0.048 for each comparison). In contrast, the ovaries of 6-week old mice in the HFD condition did not differ from the ovaries of 8-week old mice in the NC condition or 8-week old mice in the HFD condition (all adjusted *p* ≥ 0.17; Fig. 4b). Thus, exposure to HFDs at 6 weeks reduced follicle size to values comparable to those observed in older (8-week) ovaries, consistent with an accelerated progression toward a more “mature” follicle size distribution. In parallel, we quantified seminiferous tubule diameter in the testes as a morphological marker of sexual maturation in males (see methods). Males in the HFD condition showed trends towards enlarged seminiferous tubules at 6 weeks, compared to males in the NC condition, similar to the size observed in 8-week animals (Fig. 4b, Supplementary Fig. 4c). However, this difference was not statistically significant.

**Figure 4.**
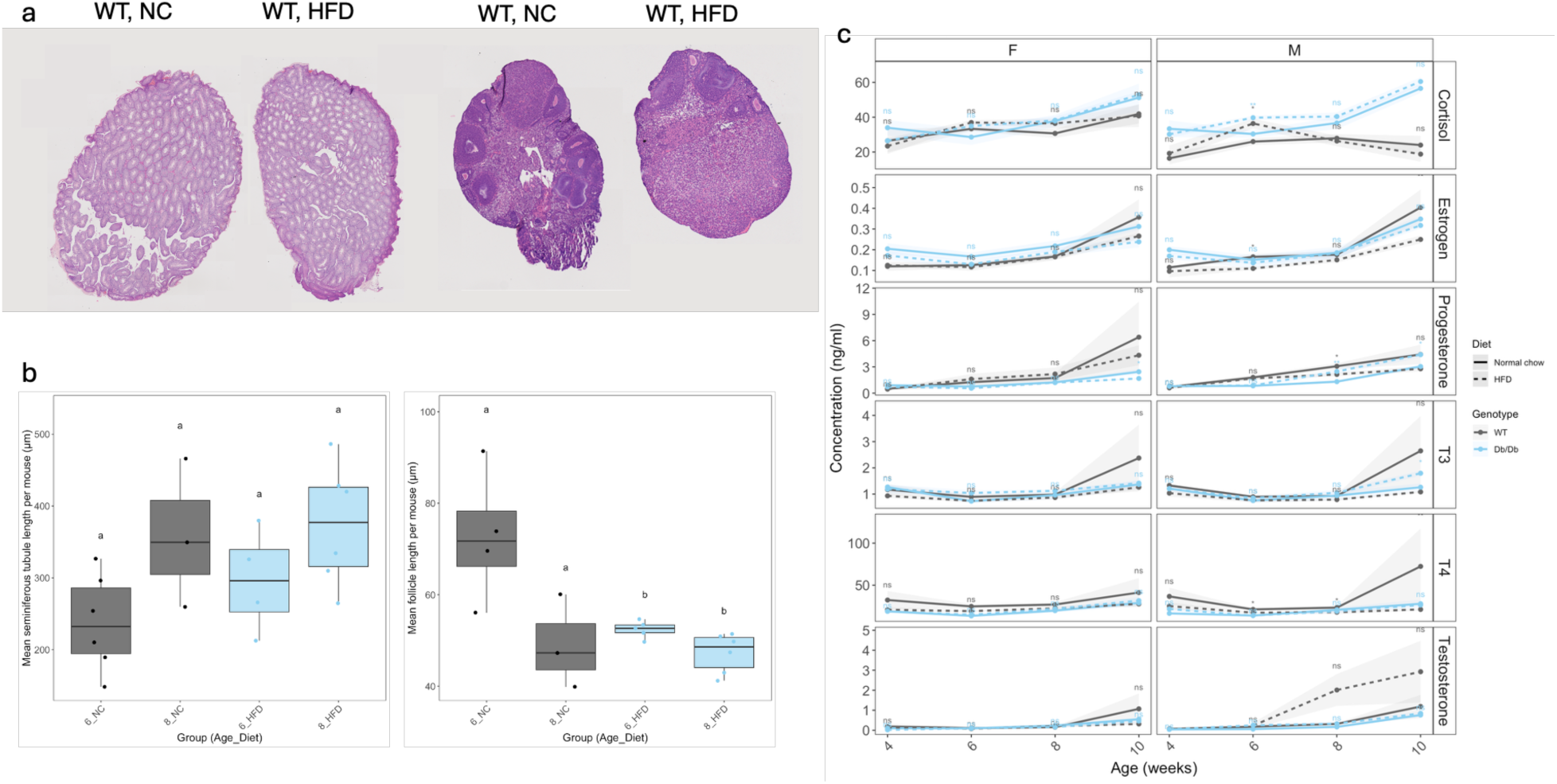
High fat diets alter the maturation of gonadal tissues and concentrations of endocrine hormones. **a)** Representative histological sections of testes and ovaries from 8-week-old WT mice in the NC and HFD conditions, illustrating follicle architecture and seminiferous tubules. **b)** Quantification of follicle length and seminiferous tubule diameter at 3, 6 and 8 weeks in WT mice in the NC and HFD conditions. Each point represents the mean value per animal (averaged across multiple measurements); boxplots show median and interquartile range. Dietary effects at 6 and 8 weeks were assessed using pairwise Wilcoxon rank-sum tests with Benjamini–Hochberg correction. **c)** Longitudinal plasma hormone concentrations (mean ± s.e.m.) for cortisol, estrogen, progesterone, T3, T4 and testosterone from 4 to 10 weeks in WT and *Db*/*Db* mice in the NC and HFD conditions, stratified by sex and genotype. At each age, NC and HFD were compared within each hormone × sex × genotype stratum using two-sided Wilcoxon rank-sum tests (ns *p* ≥ 0.05, \**p* < 0.05, \*\**p* < 0.01, \*\*\**p* < 0.001; *n* = 3–6 per group).

To place these tissue changes in an endocrine context, we measured circulating cortisol, estrogen, progesterone, T3, T4 and testosterone at 4, 6, 8 and 10 weeks in WT and *Db*/*Db* mice in the NC and HFD conditions (Fig. 4c). Both WT and *Db*/*Db* males in the HFD condition displayed increased cortisol concentrations at 6 weeks. With respect to thyroid hormones and sex steroids, WT males in the HFD condition showed higher T4 and estrogen levels at 6 weeks, and elevated T4 at 8 weeks, relative to mice in the NC condition. In contrast, hormonal profiles in the females did not respond to the diet at any time point. These results suggest that obesogenic diets elicit a sex-differentiated endocrine profile, with WT males displaying elevated HPA-axis and thyroid activity response to excess energy availability. In contrast, female systemic hormone levels remain comparatively unaffected despite clear diet-induced shifts in gonadal histology.

To understand how HFDs might regulate how energy is allocated between growth (skeleton) and reproduction (gonadal tissues), we assessed insulin receptor expression in the growth plates and gonadal tissues of animals in the HFD condition compared to the NC condition. Specifically, we reasoned that elevated insulin receptor expression could increase glucose uptake by cells to enable rapid division. We performed immunohistochemistry on histological sections taken from 6-, 8- and 10-week old animals. We observed trends towards decreased insulin receptor expression in the HFD condition in both males and females (Fig. 5a and Supplementary Fig. 5a), which indicates earlier terminal growth plate activity and potentially earlier cessation of long bone growth. These trends were detectable by 8 weeks of age in both male and female mice. This could indicate that HFDs shorten the time to reach terminal long bone elongation as growth plate closure approaches (Fig. 5b and Supplementary Fig. 5b). We also assessed insulin receptor expression in gonadal tissues at 6 and 8 weeks of age (Fig. 5c). We observed trends towards higher insulin receptor expression in the HFD condition in male mice at both time points (Fig. 5d). In contrast, females showed only mild trends toward elevated receptor expression at 8 weeks of age (Fig. 5d).

**Figure 5.**
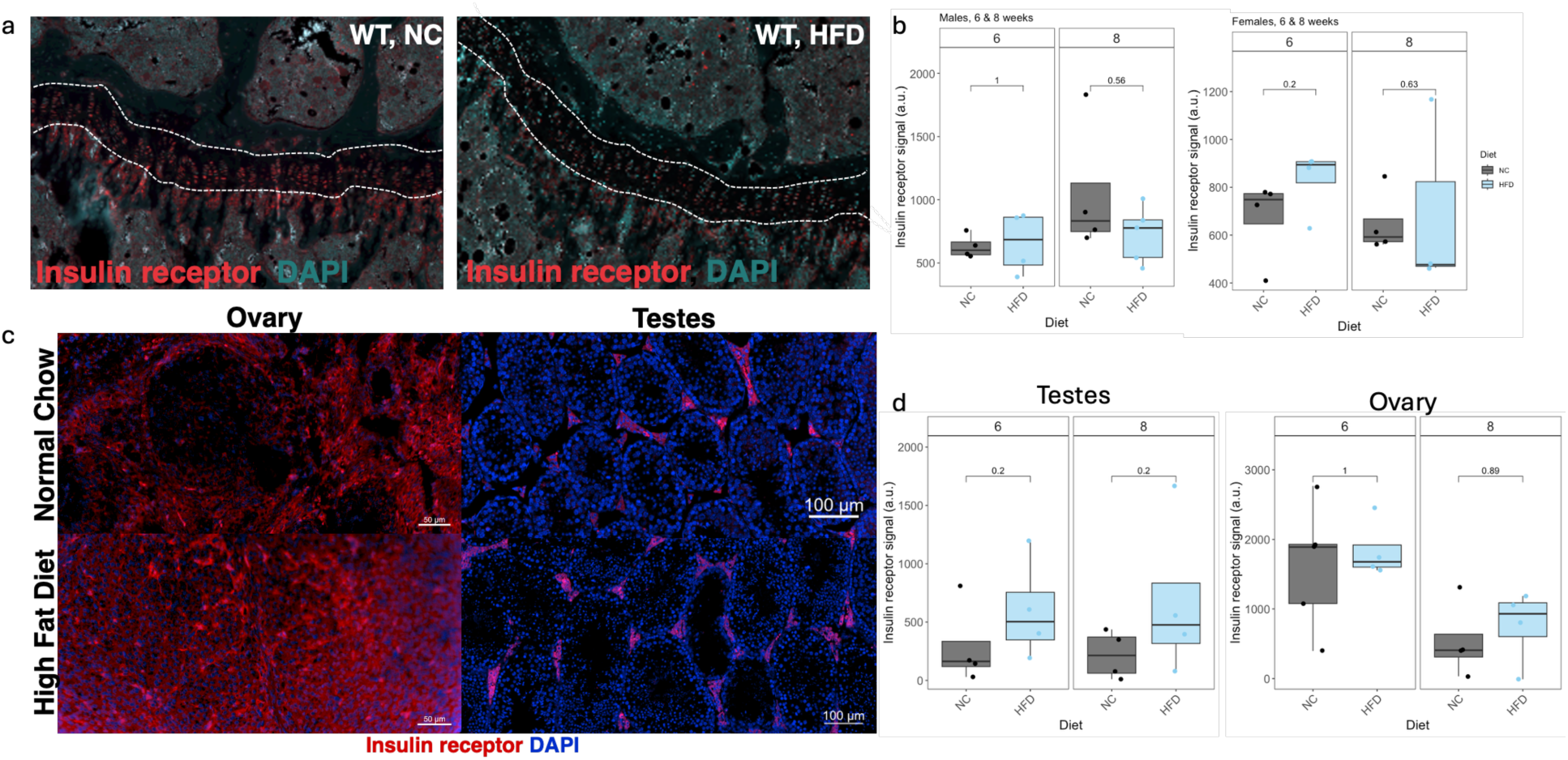
High fat diet disrupts insulin receptor expression in the growth plate and gonadal tissues. **a)** Immunofluorescence staining of growth plate cartilage chondrocytes from representative 10-week old males in the NC and HFD conditions. The curved white broken line marks the growth plate. **b)** Normalized insulin receptor signal in the growth plate in male and female WT mice in the NC and HFD conditions at 6 and 8 weeks of age. **c)** Representative immunofluorescence staining of ovary and testes tissue sections at 8 weeks of age in the NC and HFD conditions. **d)** Normalized insulin receptor signal in the ovaries and testes in male and female WT mice (respectively) in the NC and HFD conditions at 6 and 8 weeks of age. At each time point, the NC and HFD conditions were compared using two-sided Wilcoxon rank-sum tests; *n*= 3-6 in any group.

## Discussion

HFD exposure during sexual maturation consistently biased WT males towards a phenotype characterized by increased adiposity, impaired glucose tolerance, altered joint morphology, accelerated gonadal maturation, and potential disruption to insulin signaling. In contrast, WT females showed more muted systemic metabolic changes under the same dietary conditions, despite evidence of diet-responsive maturation trajectories in the growth plate and gonadal tissues. In *Db*/*Db* mice, which experience extreme energetic surplus in the absence of intact leptin signaling, many of these diet-mediated effects were blunted or qualitatively altered. This outcome suggests that canonical adiposity signaling through the leptin receptor is an important component of how organisms generally translate nutritional environments into coordinated developmental trajectories^30,31^.

From a life history perspective, adolescence is a developmental stage during which finite energy is reallocated from predominantly linear growth and structural maintenance toward maturation of the reproductive system12,13. Under this framework, the timing of puberty is a plastic trait that tracks energetic conditions12,14. Declines in the age at pubertal onset and menarche in high-income populations1,2,10,14, accompanied by rising childhood obesity,3,4 and global meta-analytic evidence for earlier breast development in girls,2 provide convergent evidence for the possibility that contemporary children are developing under sustained energetic surplus. The current work described here provides an experimental account of these relationships: HFD-fed animals, especially WT males, accumulate excess adiposity, exhibit histological and endocrine features of more advanced gonadal maturation, and show reduced cortical bone density and altered joint morphology without increases in limb length. In other words, additional energy is not uniformly scaling up all aspects of growth. Rather, energy is re-allocated to prioritize reproductive readiness at the expense of skeletal and metabolic health.

Our insulin receptor results provide a potential mechanistic link between systemic energy balance and these energy allocation alternatives. Insulin and related growth-factor pathways are deeply conserved regulators of cellular glucose uptake, proliferation and differentiation, and are thus natural candidates for mediating the competition for metabolic resources between tissues. In growth plates, we observed HFD-associated decreases in insulin receptor signal by IHC in both sexes by 8–10 weeks, consistent with earlier recorded shortening of proliferative zone lengths and a reduced window for longitudinal bone growth. In gonadal tissues, by contrast, HFD males showed an elevated insulin receptor signal at 6 and 8 weeks, whereas females displayed only mild increases. This suggests that under HFDs, proliferative capacity may be curtailed earlier in bone while being maintained or enhanced in gonads, biasing local energy allocation toward reproductive tissues. In *Db*/*Db* mice, these receptor-level changes were attenuated or reorganized, underscoring the interaction between leptin and insulin signaling in governing tissue-specific access to metabolic energy.

Our experimental findings parallel recently published human data on endocrine mediators of adolescent energy allocation. Longitudinal work in Mayan girls^32^ shows that higher cortisol (HPA-axis activation) is positively associated with c-peptide (a proxy for insulin secretion and potential glucose uptake) and adiponectin during puberty. Moreover, the patterns of energy uptake and storage differ pre- versus post-menarche^32^. These changes were interpreted as shifts in how metabolic energy is mobilized and stored as girls move from a phase dominated by somatic growth and maturation of the hypothalamic–pituitary–gonadal axis to one increasingly oriented toward menstrual regulation and building energy reserves for future reproduction. Our results complement and extend this by showing that energy rich conditions can activate these regulators of puberty earlier and more strongly, particularly in males, with measurable consequences for skeletal and metabolic properties.

Metabolic allocation models provide a quantitative framework for these patterns. Dynamic energy budget theory15 and recent formulations of life history theory that emphasize metabolism^33^ treat assimilated energy as the currency of fitness and impose mass–energy balance constraints on how it is partitioned among survival, growth and reproduction. Our data are consistent with a shift toward a “fast” solution under HFD conditions: WT males use increased energy to increase mass (particularly adipose tissue) and advance gonadal maturation, with the cost of reduced cortical bone fraction and impaired glucose homeostasis. WT females show subtler rebalancing, with diet-induced changes in growth plate structure, bone density, and ovarian maturation but relatively preserved systemic glucose handling. In males, HFDs engage multiple endocrine axes, HPA (cortisol), thyroid (T4) and sex steroids, consistent with a coordinated “fast” life-history strategy entailing higher metabolic rate, earlier gonadal maturation, and greater metabolic dysfunction. In females, the same diet leaves systemic axes comparatively unchanged, while local tissues in the skeleton and gonads respond comparatively more subtly. *Db*/*Db* mice, with intact energy intake, but lacking leptin feedback signaling, illustrate how disrupting a central energy-sensing channel reshapes these solutions across tissues. In all cases, insulin receptor expression patterns in bone and gonads emerge as one potential mechanism by which these evolutionary energy allocation rules are applied.

These sex differences align with evidence that female rodents often exhibit relative resilience to early HFD-induced metabolic dysfunction despite equal or greater adiposity,16–19 and with life history arguments that female fitness is tightly constrained by offspring viability and the integrity of somatic capital across repeated reproductive events.12–14 In our data, WT males expressed the clearest full trade-off: greater adiposity and leptin, impaired glucose tolerance, more mature testicular histology, cortical bone deficits and a pronounced redistribution of insulin receptor signaling from growth plates to testes. WT females exhibited a more localized rebalancing: expansion of the non-hypertrophic growth plate zone, reduced cortical BV/TV and accelerated ovarian follicle maturation, but little systemic metabolic deterioration. The same underlying problem of how to allocate finite metabolic energy to growth versus reproduction, is thus solved differently in males and females, with consequences for how obesogenic environments manifest as “precocious” phenotypes across sexes.

These findings should be considered in the context of several limitations. First, the high fat diet used here, though typical for experimental murine work, is nevertheless extreme (60% fat), and thus not truly representative of western diets. The effects of variations in this diet, including high sugar diets, diets with lower levels of fat, and “cafeteria” diets should be examined. Furthermore, we only measured circulating hormones and insulin receptor abundance but did not investigate downstream signaling (e.g. AKT/ERK activation, GLUT localization). Thus, the inferred mechanisms remain incompletely described. In addition, the *Db*/*Db* mouse model of obesity is a severe, monogenic obesity model with widespread developmental and metabolic abnormalities. As such, extrapolation of results in these mice to polygenic human obesity should be made with caution. Finally, the treatment window was brief, and we do not have longer longitudinal follow-ups to examine how these phenotypic changes impact long-term health. For example, articular cartilage was assessed only at 10 weeks, and later onset of degenerative changes could have been missed.

Nonetheless, these experimental findings intersect with contemporary concerns about earlier puberty in humans. Historical declines in age at menarche from 16-17 years to 12 years have been followed by further downward shifts in breast development and other pubertal markers^7,34^, coincident with energy-rich diets and sedentary lifestyles. Large-scale genetic and Mendelian-randomization analyses indicate that earlier puberty contributes causally to increased risks of breast and endometrial cancer and cardiometabolic disease in adulthood, beyond the effects of BMI alone ^35,36^. Our data provide causal support for the hypothesis that an obesogenic diet, confined to a defined peri-pubertal window, can alter gonadal development and hormonal secretion, reorganize insulin receptor signaling in bone and gonads, and impair skeletal properties and glucose tolerance, particularly in males. Framed within life history and metabolic theories ^17,18,20^, this suggests that modern nutritional environments may not only be shifting body size distributions but actively re-timing and re-wiring the energy allocation strategies that underlie human life histories.

## Author contributions

CRC, TDC, AS, VAT, DL, conceived the project and designed the research studies. Data collection was performed by CRC, AM, AS, EMV, LDS, GS, ASO, GR, DK, SA, JS, TK, DSG, CAB, RC. Analysis and interpretation of the data was conducted by CRC, AS, NGOI, TDC and RC. CRC, AS, and TDC composed the manuscript. CRC and TDC managed the funding source for the research project. All the authors read, edited and approved the final version of the manuscript.

## Acknowledgements

We are grateful for advice and guidance provided by Kris Sabbi in the Department of Human Evolutionary Biology. We are grateful for the Histology & Histomorphometry Services offered by the Center of Skeletal Research in Boston, MA and Histology & Histomorphometry Services offered by Harvard University. This work was funded by the Harvard Dean’s Competitive Fund for Promising Scholarship.

