## Supplementary Data for "Evolutionary trade-offs between growth and reproduction under obesogenic conditions: sex-biased skeletal and gonadal maturation in mice"

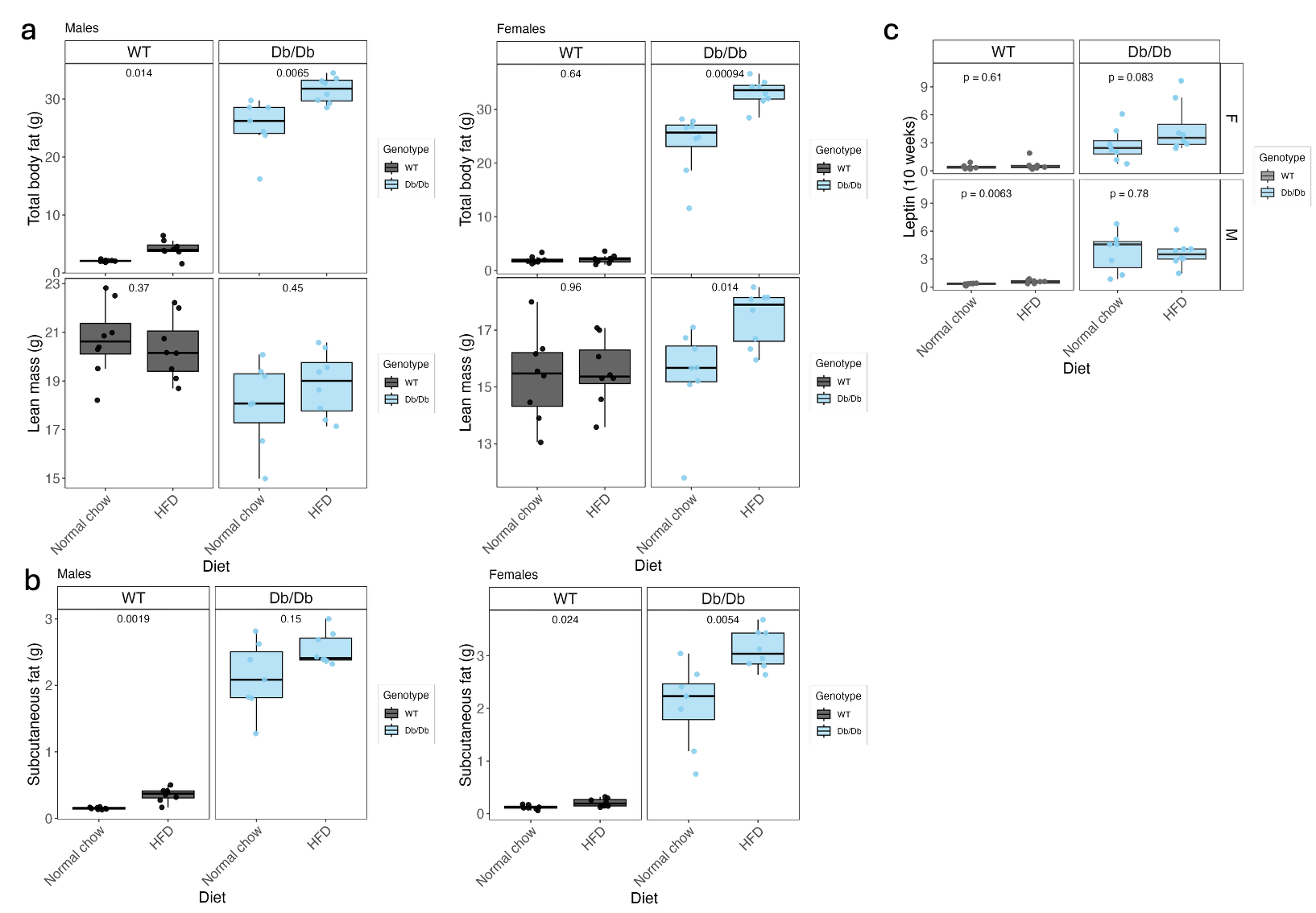

**Supplementary Figure 1. a)** Raw body fat and lean mass for male and female mice assessed by EchoMRI. **b)** Subcutaneous body fat mass at sacrifice for male and female mice. **c)** Plasma leptin concentrations (pg/ml) at 10 weeks of age for male and female mice. Wilcoxon rank–sum test (Mann–Whitney U), for each sex WT NC n = 8, WT HFD n = 8, Db/Db NC n = 7-8, Db/Db HFD n = 8.

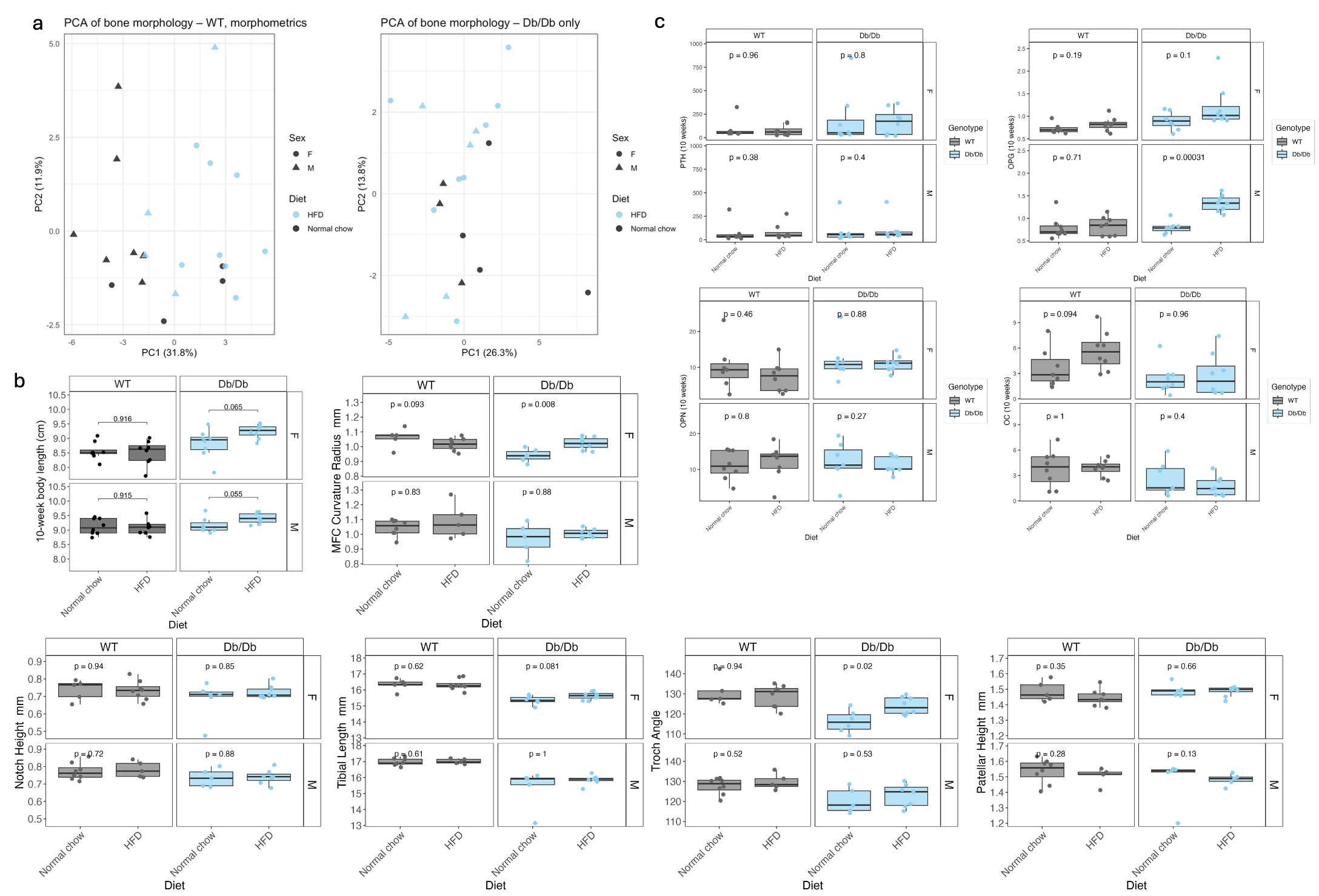

**Supplementary Figure 2. a)** PCA plots of all morphological measurements with WT only on the left and Db/Db on the right **c)** Plots of key morphological measurements that showed statistically significant differences between diets for each sex and genotype including BV/TV for cortical bone and body length at 10 weeks of age. **d)** Bone turnover markers OC, OPN, OPG and PTH. Symbols represent individual mice. Wilcoxon rank–sum test (Mann–Whitney U)). For each sex and genotype n= 5-8 in any group.

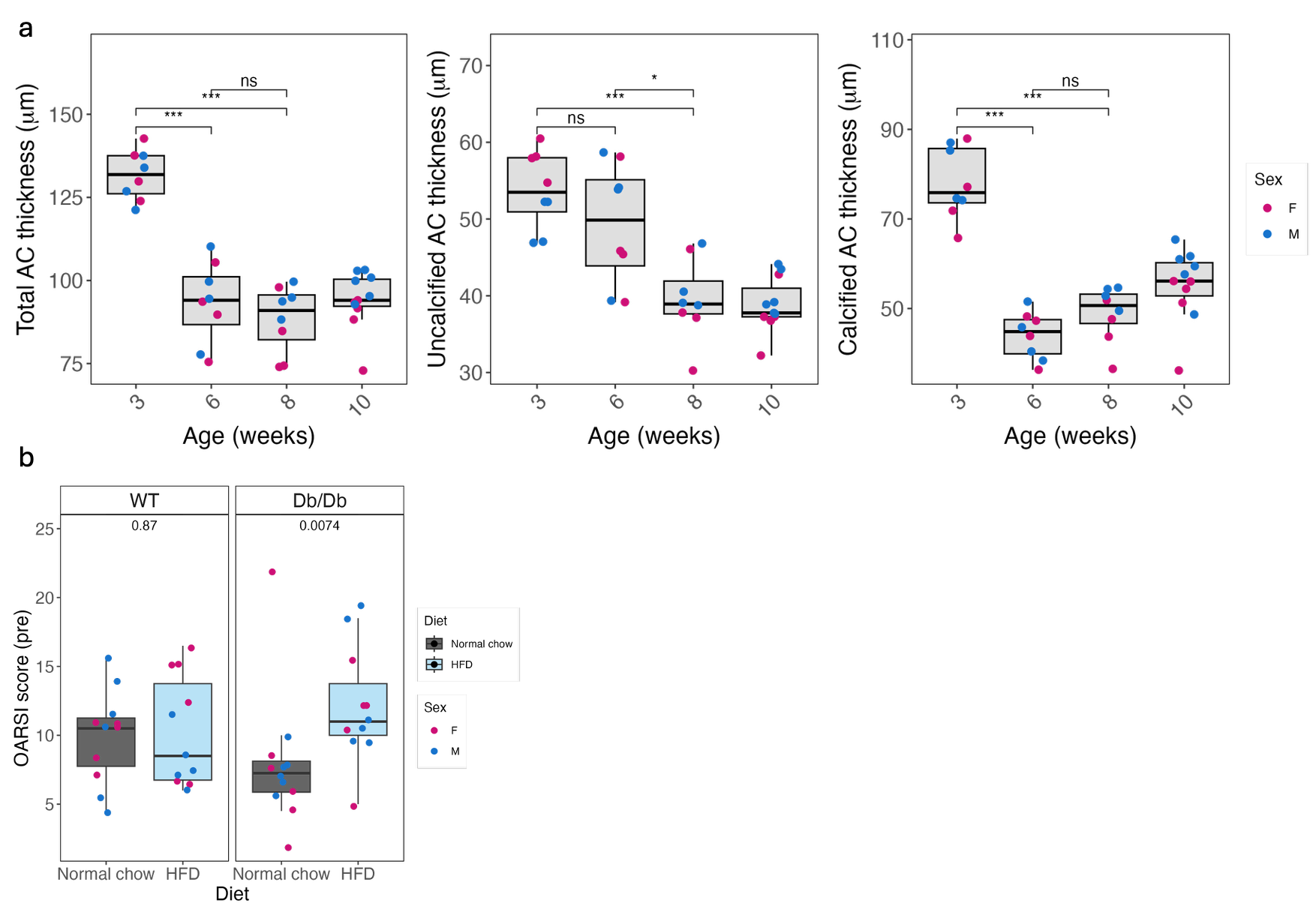

**Supplementary Figure 3. a)** WT cartilage thickness measurements of total length, uncalcified and calcified with between 3 and 10 weeks of age. **b)** OARSI scoring of articular cartilage. Summed modified Osteoarthritis Research Society International (OARSI) scores were used to assess joint damage in articular cartilage sections from 10 week old WT and Db/Db male and female animals. **Two‑sided Wilcoxon rank–sum tests** (Mann–Whitney U), for n = 8-11 in any group by age.

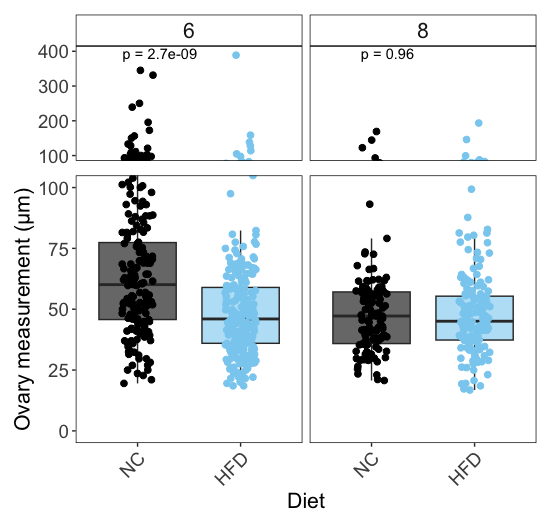

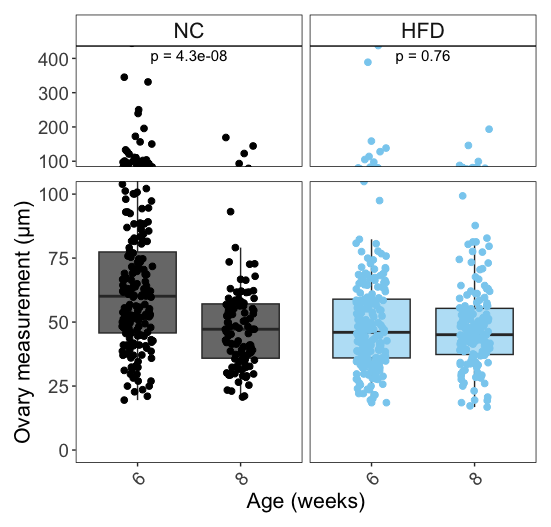

b

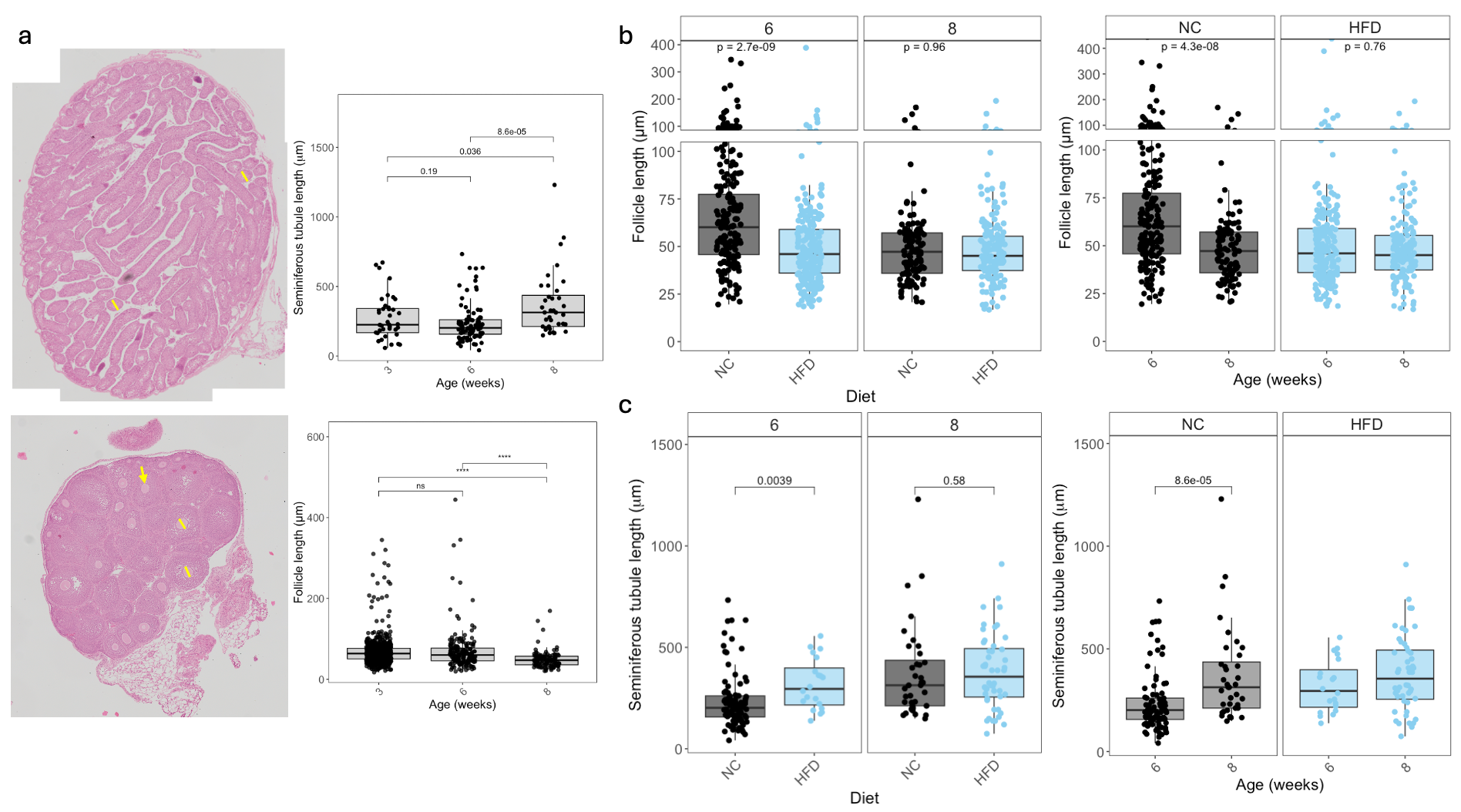

**Supplementary Figure 4. a)** Histological images of testes and ovary at 3 weeks of age accompanied by individual measurements taken of seminiferous tubules and follicles indicated by yellow lines as an example of measurements taken. Yellow arrow indicates follicle. **b)** All follicle length measurements separated by age and diet **c)** All seminiferous tubule length measurements separated by age and diet. Kruskal- Wallis test used to compare mean follicle/seminiferous across the groups (6‑week NC, 6‑week HFD, 8‑week NC, 8‑week HFD).

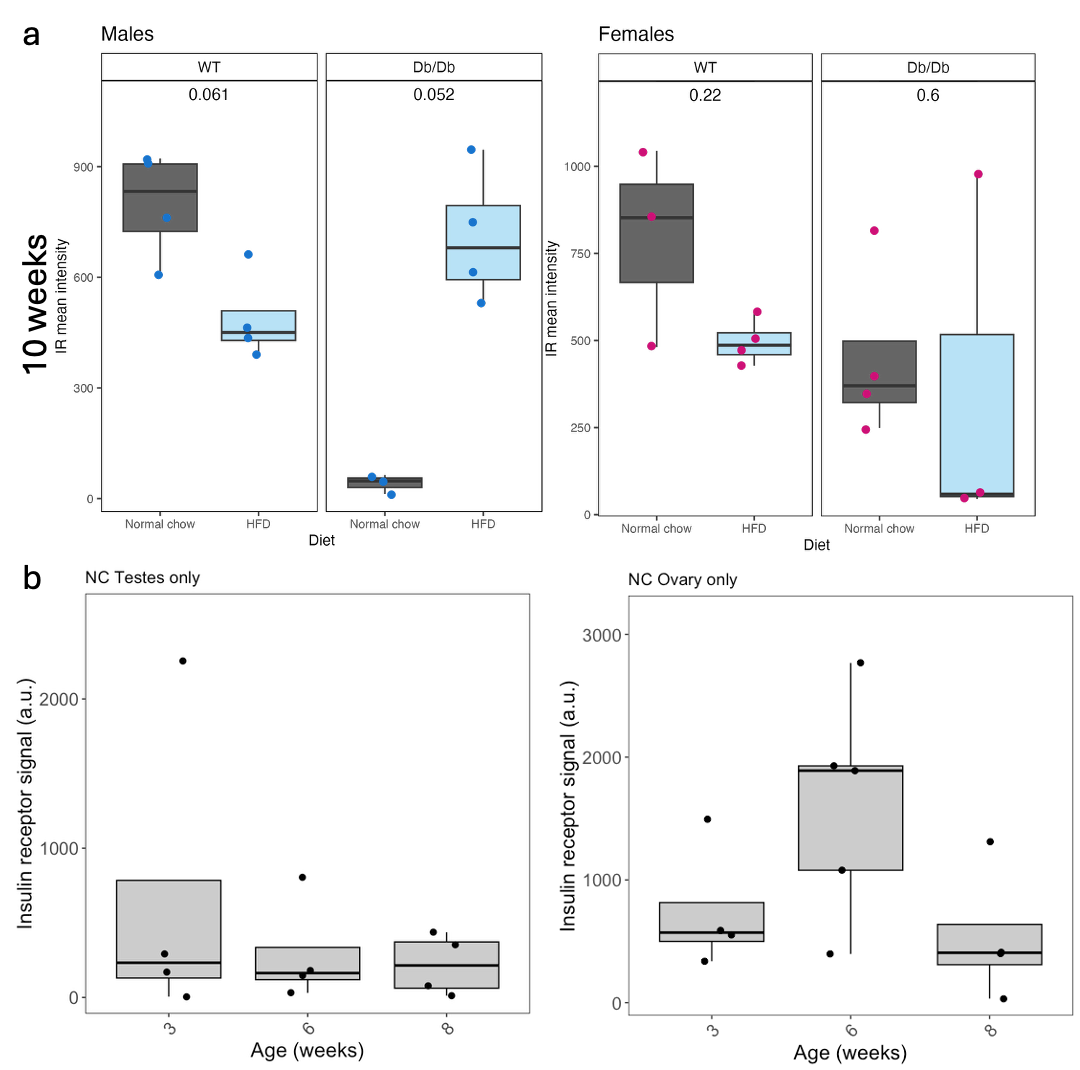

**Supplementary Figure 5. a)** Normalized insulin receptor signal in the growth plate in male and female WT mice on a NC and HFD at 10 weeks of age **b)** Normalized insulin receptor signal of WT ovary and testes tissue sections at 3, 6 and 8 weeks of age on a normal chow or high fat diet. At each age, NC and HFD were compared using two‑sided Wilcoxon rank‑sum tests; n= 3-6 in any group.
